# The use of a chimeric rhodopsin vector for the detection of new proteorhodopsins based on color

**DOI:** 10.1101/162016

**Authors:** Alina Pushkarev, Gur Hevroni, Sheila Roitman, Jin-gon Shim, Ahreum Choi, Kwang-Hwan Jung, Oded Béjà

## Abstract

Student microbial ecology laboratory courses are often conducted as condensed courses in which theory and wet lab work are combined in a very intensive short time period. In last decades, the study of marine microbial ecology is increasingly reliant on molecular-based methods, and as a result many of the research projects conducted in such courses require sequencing that is often not available on site and may take more time than a typical course allows. In this work, we describe a protocol combining molecular and functional methods for analyzing proteorhodopsins (PRs), with visible results in only 4-5 days, that do not rely on sequencing. PRs were discovered in oceanic surface waters two decades ago, and have since been observed in different marine environments and diverse taxa, including the abundant alphaproteobacterial SAR11 group. PR subgroups are currently known to absorb green and blue light, and their distribution was previously explained by prevailing light conditions - green pigments at the surface and blue pigments in deeper waters, as blue light travels deeper in the water column. To detect PR in environmental samples, we created a chimeric plasmid suitable for direct expression of PRs using PCR amplification and functional analysis in *Escherichia coli* cells. Using this assay, we discovered several exceptional cases of PRs whose phenotypes differed from those predicted based on sequence only, including a previously undescribed yellow-light absorbing PRs. We applied this assay in two 10-days marine microbiology courses and found it to greatly enhance students’ laboratory experience, enabling them to gain rapid visual feedback and colorful reward for their work. Furthermore we expect this assay to promote the use of functional assays for the discovery of new rhodopsin variants.

## Introduction

Microbial retinal-based ion pumps were first discovered in the hypersaline dwelling archaea *Halobacterium salinarum* (Oesterhelt and Stoeckenius, 1971). Since then, rhodopsins have been found in various microorganisms, spanning the three domains of the tree of life (Béjà et al., 2013; Pinhassi et al., 2016), and were even detected in viruses (Yutin and Koonin, 2012; Philosof and Béjà, 2013).

The first bacterial rhodopsin was discovered in the abundant uncultured proteobacterial SAR86 group, and was therefore named proteo-rhodopsin (PR) (Béjà et al., 2000). PRs are light-driven proton pumps that absorb light in the blue or green regions of the visible light spectrum according to the light available at the depth from which they are isolated (Béjà et al., 2001). The dominant residue responsible for spectral tuning in PRs resides in the retinal-binding pocket at position 105, with leucine or methionine in green-absorbing. PRs (GPRs) and glutamine in blue-absorbing PRs (BPRs) (Man et al., 2003; Gómez-Consarnau et al., 2007). PRs are abundant in the marine environment, and a recent metagenomics survey estimated that on average, over 60% of small microbial cells in the photic zone carry rhodopsin genes (Finkel et al., 2012).

The search for novel rhodopsins is based mostly on sequence homology screens utilizing metagenomics data collected from various environments (Venter et al., 2004; Sabehi et al., 2005; Rusch et al., 2007), or PCR performed on environmental DNA samples using degenerate primers designed for conserved regions in microbial rhodopsin proteins (Atamna-Ismaeel et al., 2008; Sharma et al., 2009; Koh et al., 2010). Currently, there are only two functional screens to search for new rhodopsins, (*i*) colony color by plating fosmid libraries on retinal containing plates (Martínez et al., 2007); and (*ii*) pH changes of fosmid clones in response to illumination (Pushkarev and Béjà, 2016).

In order to combine sequence homology and function-based methods, we devised a protocol based on a previously designed chimeric PR construct (Supplementary Fig. S1). This chimeric construct was used to express individual partial PR sequences recovered from the environment via PCR amplification, cloning and sequencing (Choi et al., 2013). Here, we improved the chimeric PR construct to enable the screening of diverse partial PR sequences directly from the environment, enabling rapid visualization of PR activity. In this manner, we developed a simple way to demonstrate the concept of niche adaptation and spectral tuning to undergraduate and graduate students. Student labs in marine microbial ecology or marine microbiology are usually offered as condensed courses ranging in length from between 10 to 30 days. Hence, there is a need for short experiments that can demonstrate some proofs of concept within the course’s timeframe.

In this work, we present a protocol (Fig. 1) for analyzing microbial samples using functional and molecular methods in short time spans, enabling students to perform high-level molecular work while receiving immediate visual results.

**Figure 1.**
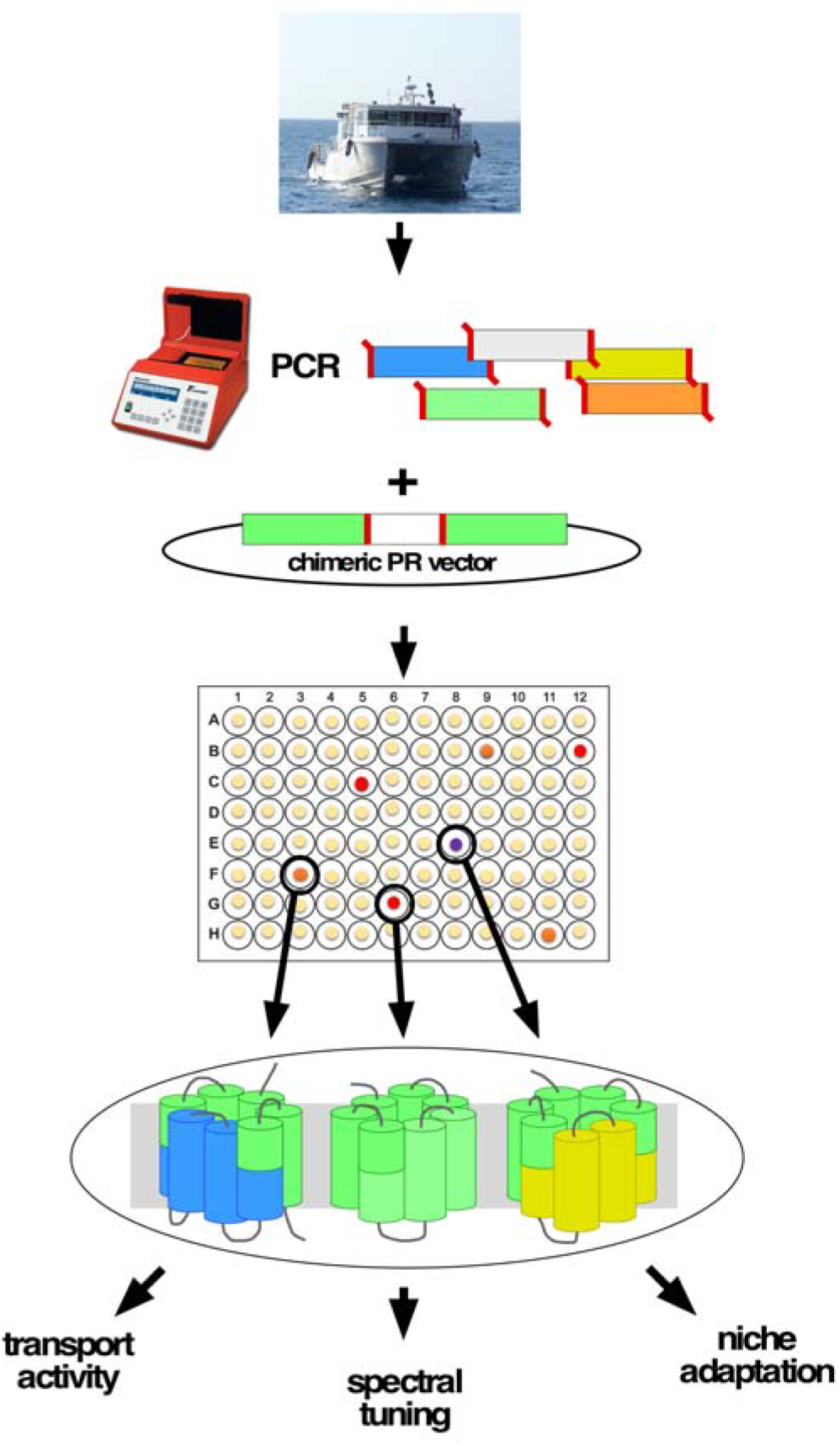
A schematic representation of the suggested protocol for the use of chimeric rhodopsin. DNA samples are obtained by various methods from the desired niches, then a PCR reaction is carried out using degenerate primers listed in Table 1. The PCR product is then digested and cloned directly onto the expression vector. The resulting colonies are harvested into individual cells in a 96-well format for storage, expression and further study.

**Table 1.**
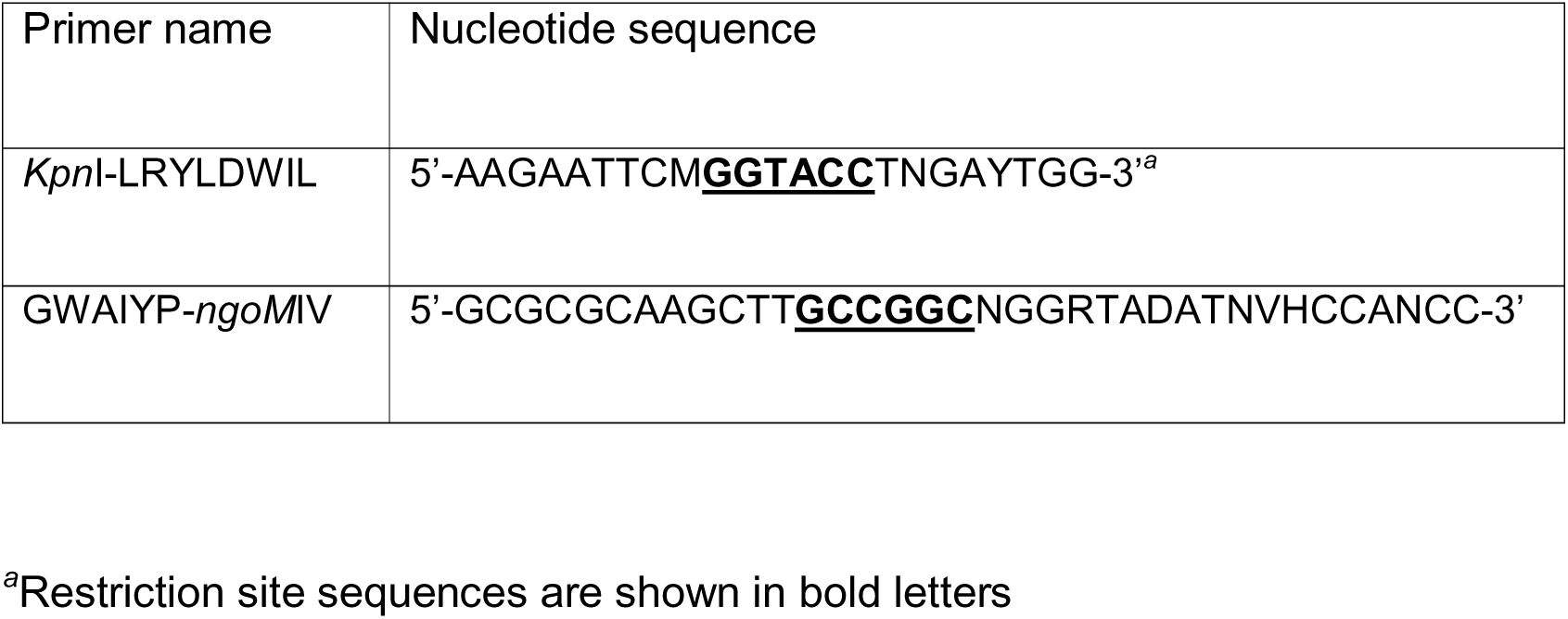
Degenerate primers designed to amplify diverse PRs using PCR reactions.

## Materials and Equipment

### Environmental sampling, DNA extraction and PCR

Sampling was performed in March 2014 in the Gulf of Aqaba, Station A (29°28′N, 34°55′E) (Supplementary File S1). Twenty liters of water from 0, 20, 40, 60, 80 and 100 meters were filtered through a GFD filter (Whatman) and collected on two 0.22 *μ*m Durapore filters (Millipore). DNA was extracted from both filters using a phenol-chloroform protocol (Wright et al., 2009).

Degenerate primers were designed based on a multiple sequence alignment of known PRs for maximum diversity coverage using the conserved regions in helixes C and F (based on a set of primers previously reported by us, (Atamna-Ismaeel et al., 2008, 2010) and modified to include *Kpn*I and *ngoM*IV sites). Each PCR reaction was performed using Takara Ex Taq^™^ polymerase (Takara-bio, Korea). Polymerase chain reaction amplification was carried out in a total volume of 50 *μ*l containing 1 *μ*l DNA template, 0.2 mM dNTPs, 1X Ex Taq buffer (Mg^2+^ plus), 0.6 *μ*M primers (each) and 125 U Takara Ex Taq^™^ DNA polymerase. The amplification conditions included steps at 98°C for 5 sec, 40 cycles of 98°C for 10 sec, 50°C for 1.5 min and 72°C for 2 min.

### Cloning and expression

PCR products were double-digested with *Kpn*I and *NgoM*IV in NE Buffer^™^ 1.1 buffer (New England BioLabs) for 2 hours at 37°C, cleaned using ½ volume of phenol and ½ volume of chloroform then 1 volume of chloroform, and heated to 70°C for 10 min to remove residual chloroform by evaporation.

The cloning vector was extracted from cells using only QIAprep Spin Miniprep Kit (Quiagen) since other methods do not enable efficient restriction, digested with *Kpn*I and *NgoM*IV in 1.1 buffer (New England BioLabs), and separated from the 114 bp insert on 1% agarose gel. Cut vector was extracted from gel using NucleoSpin® Gel and PCR Clean-up (Macherey-Nagel, Düren, Germany). Cloning vector and insert were ligated overnight using T4 DNA ligase (Thermo Scientific, Lithuania) at 4°C. The next day, ligation was inactivated using 70°C for 10 min and dialyzed for 1 hour on VSWP-25 filters (Millipore) against DDW. Ligations were transformed into electro-competent DH10B *E. coli* cells (4.5 μl into 30 μl cells), and shaken in 0.5 ml SOC medium at 37°C and 200 rpm. 0.25 ml was plated on LB-agar ampicillin (Amp) plates and incubated overnight. Colonies were transferred to 96-well plates with 230 μl LB-10% glycerol Amp for long-term storage. For initial color expression, plates were duplicated to U-shaped bottom 96-well plates (Thermo Scientific, Denmark) with 150 μl LB, 50 μg/ml Amp, 1.2 mM IPTG and 15 *μ*M all-*trans* retinal, shaken at 37°C and 250 rpm, and covered overnight with AeraSeal^™^ air permeable sheet (EXCEL Scientific, California, USA). Plates were centrifuged at 3,000X RCF (Sigma 4-16KS centrifuge) at room temperature for the detection of cell color.

### Absorption assay of intact cells and purified protein

A fresh colony was used to inoculate 50 ml LB 50 μg/ml Amp, 1.2 mM IPTG and 15 μM all-*trans* retinal, and shaken at 37°C and 200 rpm in a 125 ml Erlenmeyer. Cells were collected by centrifugation, washed twice with buffer A (50 mM Tris-HCl pH 8 and 5 mM MgCl_2_), then resuspended in 1 ml of the same solution. The absorbance of the supernatant was recorded between 400 and 800 nm using a spectrophotometer (Shimadzu UV-1800, Japan) against buffer A blank. In addition, absorbance spectra of purified protein was measured as described previously (Choi et al., 2013). The results are summarized in the supplementary material (Supplementary File S2) with the subtraction of the negative control signal and purified protein spectra.

### Sequencing and phylogenetic analysis

All clones were extracted using the standard alkaline lysis miniprep protocol and sequenced using standard M13R primer (GCGGATAACAATTTCACACAGG, Macrogen, Korea). The unique middle parts of each clone, excluding any primer sequences, were deposited in GenBank under accession numbers KY963379-KY963416, eliminating redundant sequences at each depth. Full detailed chimeric sequences of all clones are available in the supplementary material (Supplementary File S4).

A maximum likelihood phylogenetic tree was constructed using the phylogeny.fr pipeline (Dereeper et al., 2008), which included PhyML v3.0 (Guindon et al., 2010), the WAG substitution model for amino acids (Whelan and Goldman, 2001) and 100 bootstrap replicates.

### Proton pumping activity assay

Cells were inoculated from a fully thawed plate into two 96-well 2.2 ml plates (ABgene, UK, Cat. No. AB-0932) filled with 1 ml LB (50 μg/ml ampicillin, 1.2 mM IPTG and 15 *μ*M all-*trans* retinal) in each well and grown at 30°C, shaken at 700 rpm for 17 hours. Cells were collected by centrifugation at 3,000X RCF (Sigma 4-16KS centrifuge) and washed twice with 0.5 ml minimal salt solution (10 mM NaCl, 10 mM MgSO_4_ and 10 mM CaCl_2_). Finally, cells from both plates were re-suspended in a 150 μl (final volume) minimal salt solution in dark well plates with a transparent bottom (Greiner Bio-One, Cat. No. 655096). Cells were allowed to settle for 10 min in the dark at RT, after which a functional screening was performed using a customized robotic system (TECAN, Männedorf, Switzerland) as follows: pH from eight pH electrodes (Sentek UK, Cat. No. P13/2.5M/BNC) was recorded simultaneously by a multi-parameter analyzer (Consort, Belgium, model 3060) with readings logged every 1 sec for 3 min of dark, followed by 2 min of illumination by 2 LED lights, warm white light (2,600-3,700 CCT spanning 420-700 nm, intensity 12 lm, Cree Inc.) and blue light (485 nm peak, flux 10.7 lm, Cree Inc.), constituting all visible spectra under each well. The dark/light cycles were measured twice to confirm consistency. Eight wells were measured simultaneously, each illuminated by two LED lights at a distance of 17 mm between the LEDs and the cells, a setup constructed by Neotec Scientific Instrumentation Ltd. (Kfar Saba, Israel). Each well received a light intensity of 450 μmol photons m^-2^ s^-1^.

## Abbreviated Protocol

Please add here RV1_SUPPfileS3.docx

## Results and discussion

We used a GPR-containing vector with designed restriction sites (Choi et al., 2013) and replaced the middle part of the PR with a shorter random DNA sequence that is incompatible with the open reading frame (ORF) of the third part of the chimera (Supplementary Fig. S1), thereby introducing a premature stop codon. This new vector (pKa00X) was created to avoid the high background (reddish colonies) observed with the original chimeric GPR vector (pKa001), which does not allow distinguishing newly found GPR from an uncut vector. The chimeric rhodopsin also contains a C-terminal 6-His tag, allowing the purification of positive proteins for further characterization.

We used extracted DNA samples collected from multiple depths (0 to 100 meters at 20-meter intervals). All samples tested positive for PRs, resulting in two PCR amplicons of ~400 bp and ~330 bp (Supplementary Fig. S2). Two 96-well plates of clones were picked for each depth to detect colored colonies for further study (Fig. 2). Out of a total of 1,152 colonies, 45 had visible color and were chosen for further characterization. The PRs obtained showed absorption spectra expected from the depth they were collected from, with yellow-absorbing PR (YPR; purple colonies) and GPR (red colonies) found in surface waters, while BPRs (orange colonies) were found mostly in the deeper samples (Fig. 3A). The visual results allowed us to selectively choose the different positive phenotypes for further analysis and sequencing. The PRs amplified by our primers were diverse, with some being similar to PRs from different microbial groups ranging from Marine Group II (MGII) Euryarchaeota through the proteobacterial groups SAR11, SAR86 and SAR92, to Flavobacteria (Fig. 3B).

**Figure 2.**
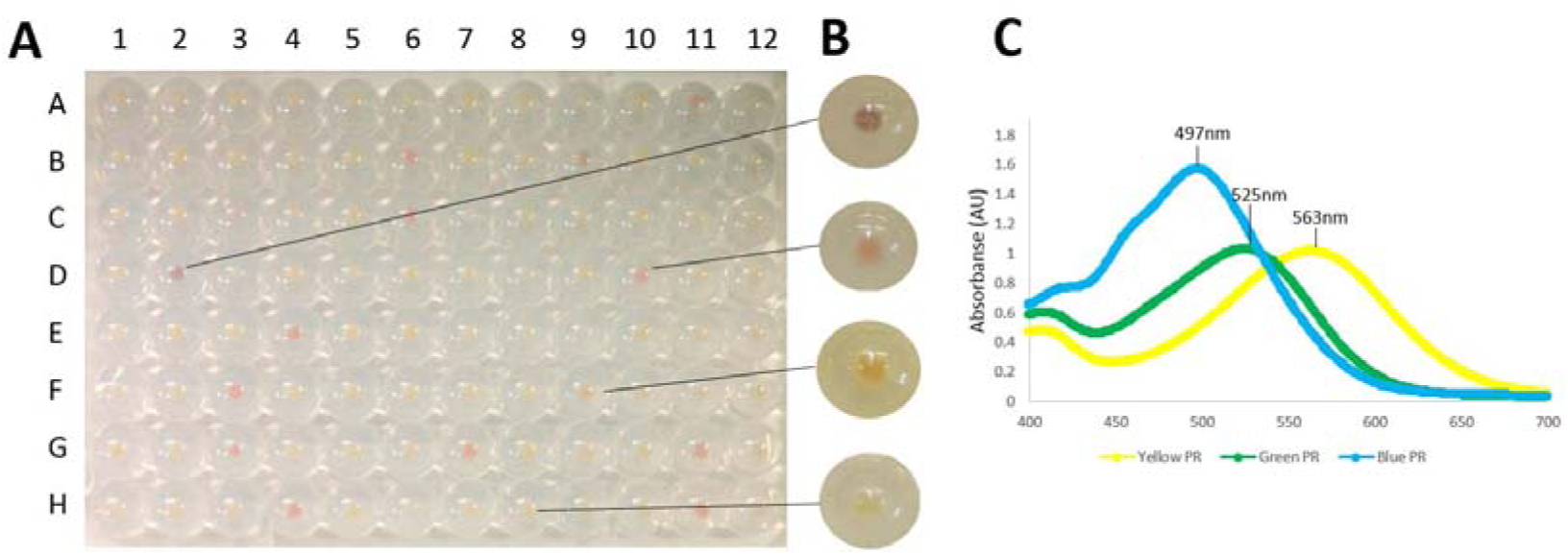
A photograph demonstrating results from one of the expression plates in this study. (A) Colonies were grown in 96-well plates as described in Materials and Methods, induced by IPTG in the presence of all-*trans* retinal. (B) A close-up of representative clones (from top to bottom): yellow-, green‐ and blue-absorbing chimeric rhodopsin, and empty vector pKa00x. (C) Absorption spectra of purified protein representing yellow-, green‐ and blue-absorbing chimeric rhodopsins (BPR; #42, YPR; #1, GPR; #11).

**Figure 3.**
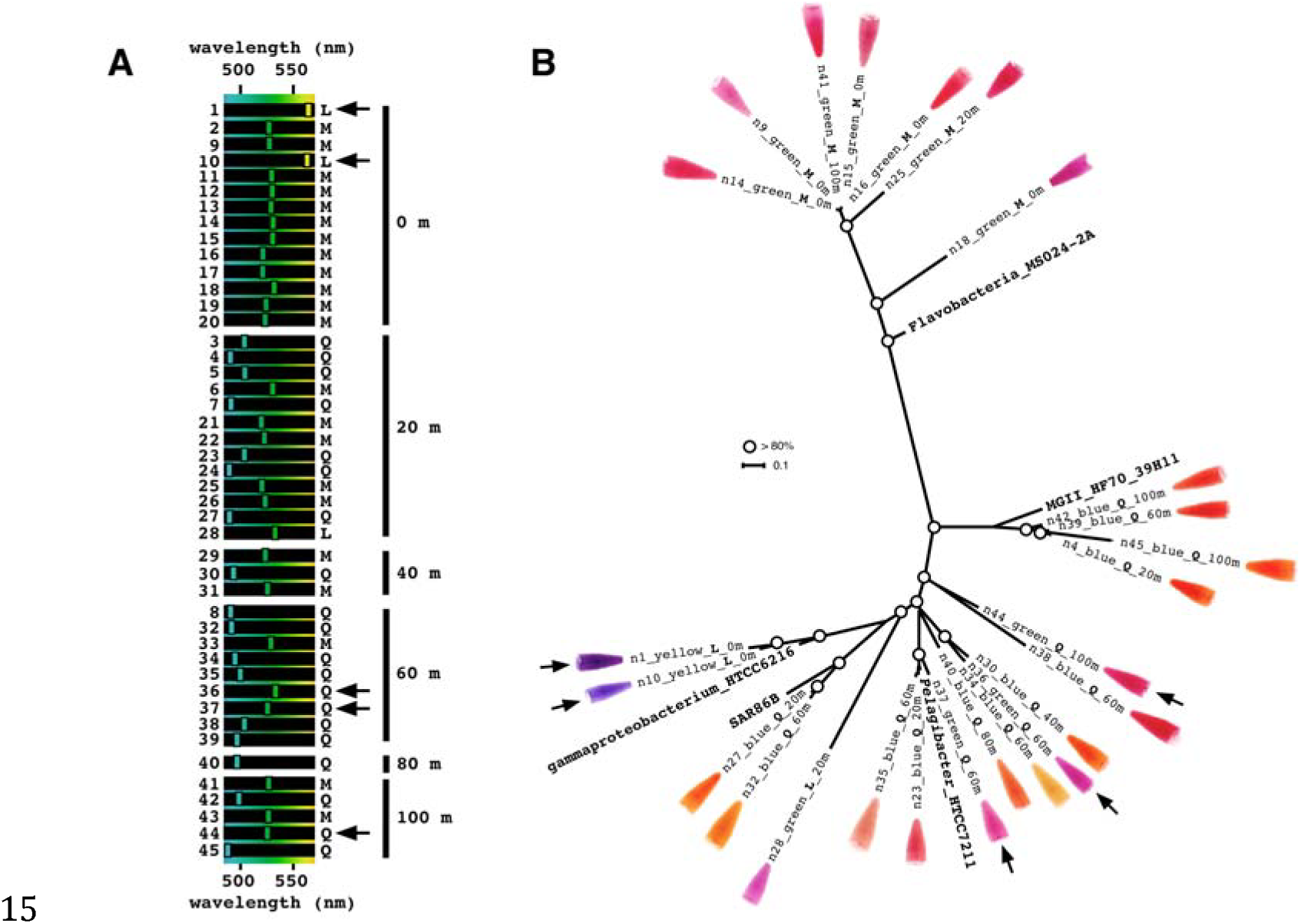
Diverse chimeric clones obtained from different depths in a single cast at station A, the Red Sea. Black arrows mark the clones showing discrepancies between genotype and phenotype according to homologous position 105. (A) A phylogenetic tree was constructed using unique representatives from each depth with rhodopsins from representative cultured microbes. Circles represent bootstrap values higher than 80%. Next to each clone appears a picture of purified protein after induction in the presence of all-*trans* retinal. (B) The absorption spectrum of each clone was measured after protein purification, and the peak is represented on the spectrum between 480 and 570 nm. Clones are arranged according to depth.

Position 105 in PR is believed to be the main determining position for PR wavelength absorption (Man et al., 2003); non-polar methionine (Gómez-Consarnau et al., 2007) or leucine at position 105 results in GPR, while the polar glutamine residue at position 105 results in BPR. Several interesting exceptional cases were observed in our screen: clone 1 and 10 are YPR although they contain leucine at position 105, while clones 36, 37 and 44 appear as GPR although having a glutamine at position 105 (Fig. 3). In the case of clones 36 and 37, we can presume that the cause of the red shift could be attributed to another similar clone (clone 34, with only one amino acid change compared to clone 36; see Fig. 4) absorbed in the blue. The change is a cysteine to a tyrosine residue in the loop region between transmembranal domains E and F. This region was previously reported to influence spectral tuning (Yoshitsugu et al., 2008). Further work with clones 1, 10 and 44 should be performed in order to understand their spectral tuning mechanism.

**Figure 4.**
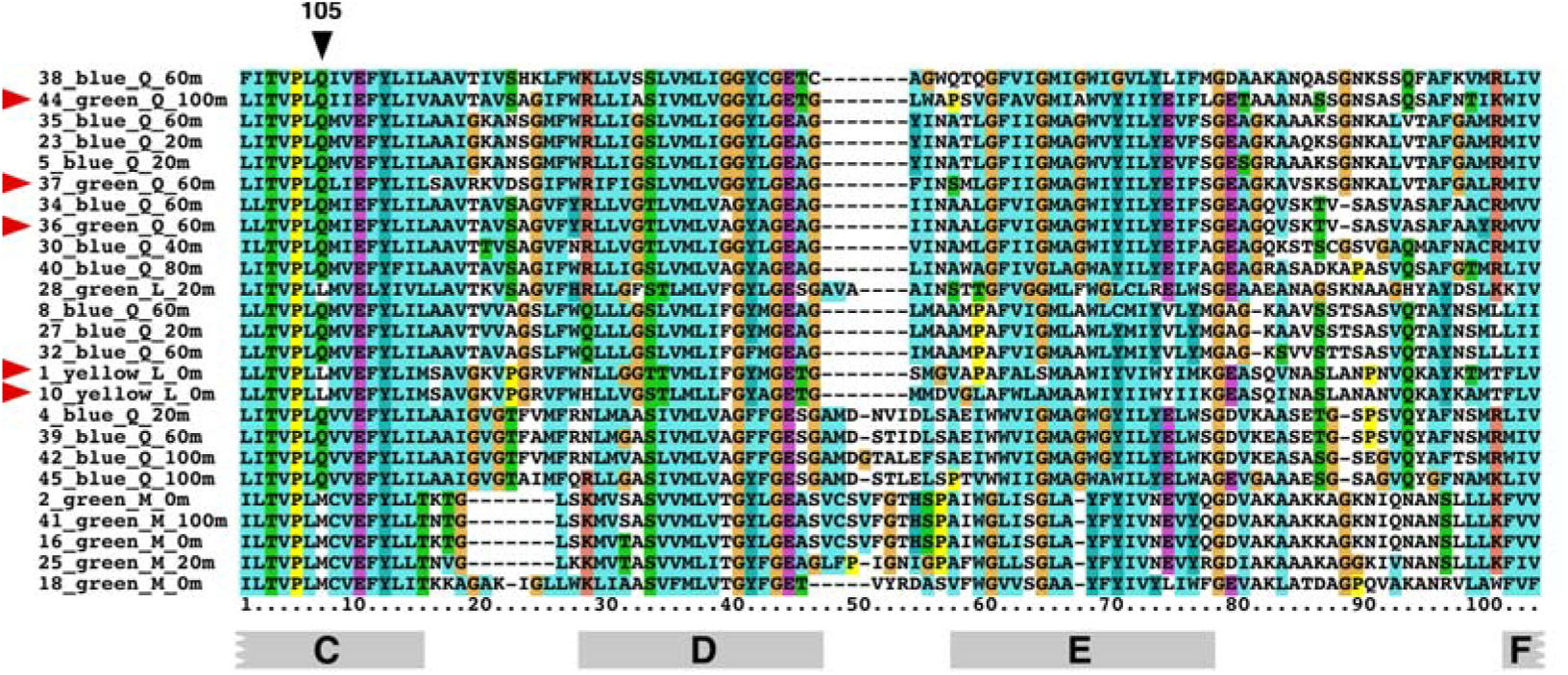
Protein sequence alignment of representative clones found in this study. Position 105 is marked (black arrow). Red arrows mark the clones showing a discrepancy between phenotype and genotype according to position 105. At the bottom of the alignment are the corresponding transmembrane helixes of PR.

Furthermore, to test the adsorption spectra, we compared our wholecell measurements with purified protein measurement (Choi et al., 2013). A detailed comparison of the two methods is presented in the supplementary material (Supplementary File S2). The advantages of whole-cell measurement are rapid results acquisition with minimal expenses and equipment, which are specifically relevant to basic laboratory courses. On the other hand, purified protein measurement has a much greater signal-to-noise ratio due to the absence of cell content interference, yet requires a longer protocol with higher costs. Both spectra obtaining methods are influenced by the pH of the medium so the exact pH should be noted at each measurement. An abbreviated and detailed protocol is attached to the supplementary material for use in teaching and research (Supplementary File S3).

### Concluding remarks

To determine if the clones are functional proton pumps, a light-driven proton pumping assay was performed as described in (Pushkarev and Béjà, 2016). Proton pumping activity was observed in all chimeric PRs with varying intensities (Supplementary File S2). This shows that although the clones generated by this assay are chimeric, they are functional (i.e. able to transport protons across the membrane), and could therefore be used for teaching and discussing various aspects connected to rhodopsins, such as the use of uncouplers, change of membrane potential, proton pumping under various wavelengths, the conversion of light energy to potential or chemical energy, and more.

Expression of environmental PR fragments depends on primer matching, correct length of PCR product, frame and compatibility to the chimeric construct. Therefore, it would be interesting to compare the chimeric ligation results to standard TA cloning of the PCR fragments. This would allow the estimation of the rhodopsins “left in the dark” in such an experiment, and a comparison of the sequences to the ones that underwent expression in the chimeric vector. This could also be useful for altering the primers to better express rhodopsins from a specific environment. Even though conserved regions between helix C and F are important for color tuning (Man et al., 2003; Choi et al., 2013), it would also be interesting to create chimeric constructs where the constant part is of BPR or YPR origin for the expression of protein and testing the dominance of each part over the spectral tuning residues.

In terms of environmental implications, the fact that we retrieved two YPRs implies that besides the known GPRs and BPRs (and now YPR), more is yet to be found. Red light penetrates as deep as 25 meters in clear oceanic water, therefore the existence of more natural YPR, as well as orange‐ and red-absorbing PR variants, are expected in future experiments with the new chimeric PR vector. Although we have tested using ocean genomic samples, this will also apply to freshwater, lakes, glaciers, etc.

## Acknowledgments

The authors would like to thank the captain and crew of the R/V Sam Rothberg of the Inter University Institute in Eilat, Israel, for their expert assistance at sea and for providing their facilities for primary processing of the samples.

This work was supported by Israel Science Foundation grant 1769/12, the I-CORE Program of the Planning and Budgeting Committee, the Grand Technion Energy Program (GTEP), The Leona M. and Harry B. Helmsley Charitable Trust reports on Alternative Energy series of the Technion – Israel Institute of Technology and Weizmann Institute of Science, and NRF grants of Korea-2016R1A6A3A 11934084 and 2015R1D1A1A01058917.

## Conflict of Interest

The authors declare no conflict of interest.

## Supplementary Material

**Supplementary Figure S1.** Chimeric construct for the expression of partial rhodopsin genes from the environment. (A) pKa001 is the original construct used for the expression of verified PR sequences in previous studies (GPR phenotype) compared to pKa00x harboring a short ORF disrupting random sequence (no phenotype) for background elimination during screening. (B) A schematic representation of the seven transmembranes of PR. The middle part is replaced by partial rhodopsins from the environment. The dominant spectral tuning residue is in the interchangeable region between helixes C and F.

**Supplementary Figure S2.** PCR resulted in PR amplification at all depths tested on March 3, 2014 at station A, the Red Sea. Primers used in this reaction are listed in Table 1. The reaction resulted in two bands of approximately 400 and 330 bp. The marker used in this 1% agarose gel is 100bp DNA Ladder RTU (GeneDirex®).

